# Automatic monitoring of whole-body neural activity in behaving *Hydra*

**DOI:** 10.1101/2023.09.22.559063

**Authors:** Alison Hanson, Raphael Reme, Noah Telerman, Wataru Yamamoto, Jean-Christophe Olivo-Marin, Thibault Lagache, Rafael Yuste

**Author notes:** Co-last authors (Thibault Lagache, Rafael Yuste).

## Abstract

The ability to record every spike from every neuron in a behaving animal is one of the holy grails of neuroscience. Here, we report coming one step closer towards this goal with the development of an end-to-end pipeline that automatically tracks and extracts calcium signals from individual neurons in the cnidarian *Hydra vulgaris*. We imaged dually labeled (nuclear tdTomato and cytoplasmic GCaMP7s) transgenic *Hydra* and developed an open-source Python platform (TraSE-IN) for the Tracking and Spike Estimation of Individual Neurons in the animal during behavior. The TraSE-IN platform comprises a series of modules that segments and tracks each nucleus over time and extracts the corresponding calcium activity in the GCaMP channel. Another series of signal processing modules allows robust prediction of individual spikes from each neuron’s calcium signal. This complete pipeline will facilitate the automatic generation and analysis of large-scale datasets of single-cell resolution neural activity in *Hydra*, and potentially other model organisms, paving the way towards deciphering the neural code of an entire animal.

## Introduction

A major goal in neuroscience is to record the activity of all neurons in a behaving animal with single - neuron resolution to capture the emergent properties of neural circuits^1^. This remains a significant technological challenge in complex systems such as rodents^2^, monkeys^3^, and humans^4^ in which whole brains can currently only be recorded with low spatiotemporal resolution. To overcome these challenges, neuroscientists have used simpler systems, including zebrafish^5–8^, *Drosophil*a^9,10^, and *C. elegans*^11,12^, which contain far fewer neurons but exhibit more limited behavioral repertoires. However, even these simpler systems are still quite complex and present their own unique imaging challenges. Here, we have chosen to focus on a small freshwater cnidarian, *Hydra vulgaris*, with one of the simplest known nervous systems^13–16^. *Hydra* has key advantages, including a transparent body, small size and number of neurons belonging to only a dozen cell types^17^, distributed nerve net lacking a true brain or ganglia, remarkable regenerative capacity^18^, and behavioral repertoire that can be automatically quantified using machine learning^19^. Unfortunately, *Hydra* is also a highly deformable organism and is therefore one of the most difficult test cases for identifying and tracking single neurons in a moving animal.

Despite much progress, monitoring of single-cell resolution whole-body neural activity in a behaving animal remains a significant challenge, even among simpler systems. Indeed, even if high-resolution imaging of whole-body neural activity can be achieved, identifying and tracking individual neurons in a moving animal over time remains a major hurdle. Moreover, the robust prediction of individual spikes from each neuron’s calcium signal also requires advanced processing of tracked calcium signals. Several approaches have been employed to facilitate the tracking of single neurons, including fixing the head of the animal to the microscope^8–10,20^, moving the microscope to keep the brain of the freely behaving animal in the center of the objective^7,11^, or immobilizing the entire animal^5,21,22^. In the first two cases, neural imaging is limited to the head only and does not allow organism-wide neural recording while behaving. In the last case, neural imaging is limited solely to immobilized, non-behaving animals. In addition to these limitations, most whole-brain imaging methods have only been achieved over short time scales with low temporal resolution.

New approaches in transgenic animals, hardware, and software have been developed to overcome these challenges. For example, a new transgenic *C. elegans* line was generated that expresses two fluorophores in the nuclei of neurons: a calcium-insensitive fluorophore (red fluorescent protein) and a calcium-sensitive fluorophore (GCaMP), which allows continuous visualization of the location of the neurons even when they are inactive^23^. These dually labeled animals make it significantly easier to detect and track individual neurons over time while the worm behaves. Novel imaging modalities include light sheet microscopy approaches, such as swept confocally aligned planar excitation (SCAPE)^24^, which has allowed fast whole-organism imaging with higher signal-to-noise and less photobleaching than widefield microscopy, and less photobleaching but poorer signal-to-noise than confocal microscopy approaches. On the software side, the advent of deep-learning algorithms has drastically improved the accuracy of single cell segmentation, a fundamental prerequisite for the tracking of neuronal activity with calcium fluorescent indicators. However, the monitoring of neuronal activity requires their robust tracking over time, which consists of pairing the neurons segmented at different time points into coherent tracks^25^. Tracking is a difficult task due to animal motion, occlusions, and intermittent activity of calcium probes. It is still performed manually or semi-automatically in many systems. Many advanced algorithms have also been developed for the accurate estimation of neuronal activity (spikes typically) from recorded calcium activity^26–30^. However, all these methods usually require signal pre-processing to attenuate imaging noise and remove non-linear fluorescence baseline that could bias the estimation of neurons’ spikes from fluorescent calcium traces. Therefore, there is a need for a robust and versatile tool that can track individual neurons in entire deforming bodies, accurately estimate the neuronal activity from calcium fluorescence, and could also be applied to more simple cases where animals are fixed or minimally moving.

Here, we developed TraSE-IN, an end-to-end pipeline (Figure 1) that allows the automatic tracking and signal extraction of individual neurons in whole behaving *Hydra*. This novel pipeline entails both hardware and software improvements over existing tools to create a versatile, robust, and accurate way to automatically detect and track individual neurons in a highly deformable organism while behaving. These improvements include the generation of a two-color transgenic *Hydra* and imaging method that allows the simultaneous recording of both nuclei and neural activity, and the development of an open-source Python platform (https://github.com/raphaelreme/trase-in). This platform consists of a tracking module to monitor the calcium activity of all neurons in the behaving animal, and a series of signal processing modules for the robust extraction of individual spikes from calcium traces, paving the way for the analysis of the functional correlates underpinning animal behavior.

**Figure 1.**
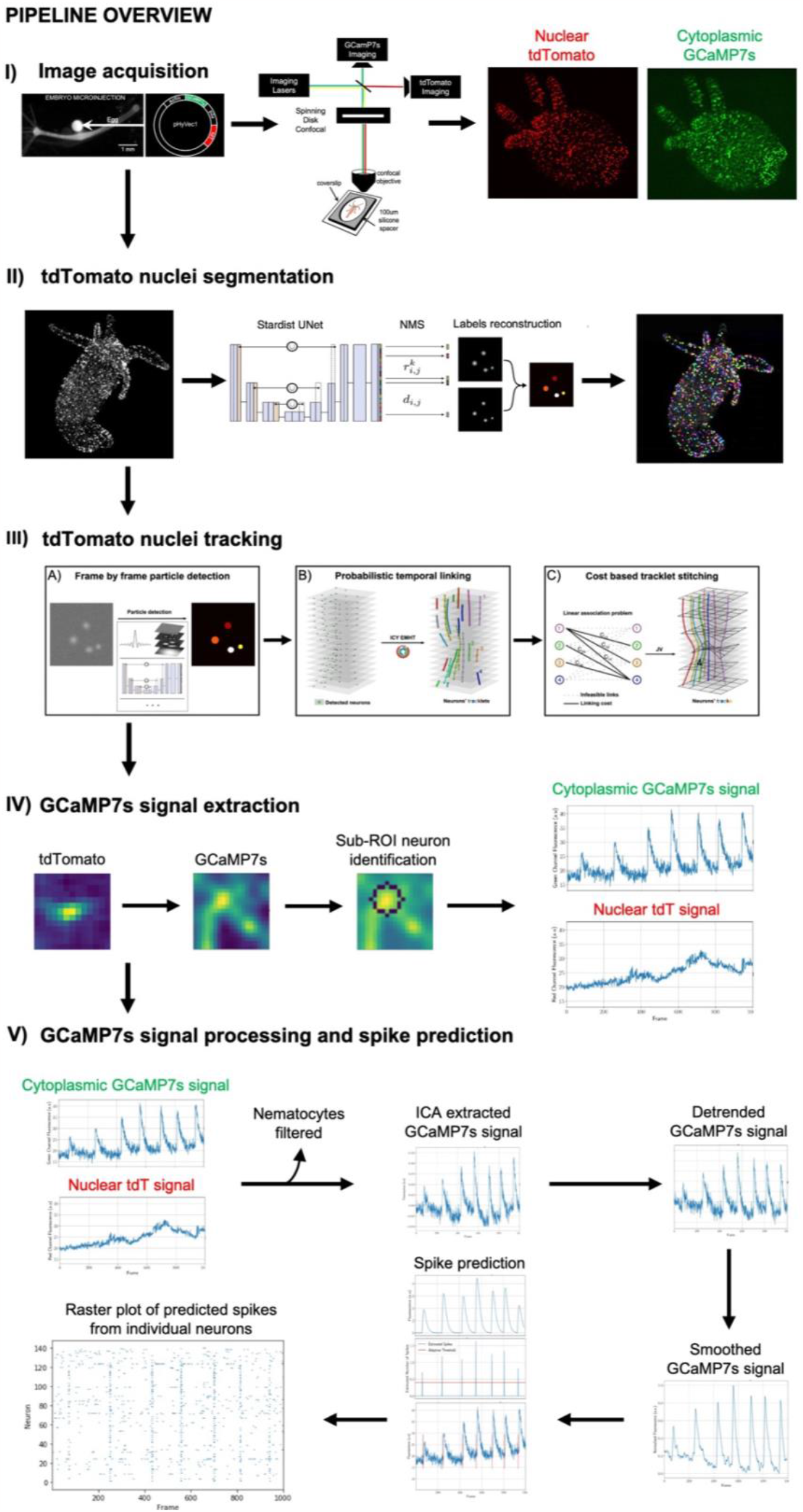
Overview of the pipeline. (I) The pipeline includes generation of a novel dually labeled (nuclear tdTomato and cytoplasmic GCaMP7s) transgenic *Hydra* imaged on a custom built simultaneous two-color spinning disc confocal microscope, allowing high spatiotemporal resolution imaging of behaving animals. (II) Each nucleus is segmented using the Stardist deep learning algorithm. (III) Each segmented nucleus is then tracked over each frame in the movie by first using probabilistic temporal linking to form short tracklets, which are then stitched together using a cost-based method to determine the final full-length tracks. (IV) A region of interest (ROI) is then drawn around the center position of each nucleus and a sub-ROI method is used to detect the most likely corresponding neuronal cell body from which to extract GCaMP7s signal. (V) Both the extracted nuclear tdTomato and cytoplasmic GCaMP7s signals are used for signal processing to account for motion artifacts. Independent component analysis, followed by detrending and smoothing is run to remove imaging artifacts from the GCaMP7s signal. Neurons’ spikes are predicted from the pre-processed calcium signal using FOOPSI. The final output is a raster plot with the spike predictions from all individual neurons in a behaving animal. See text for further details.

Altogether, hardware and software developed in the pipeline will significantly enhance our ability to correlate entire organism neural activity with behavior under different conditions, analyze large data sets over longer time scales, and bring us closer towards the goal of cracking the neural code of an entire animal.

## Results

### I. Imaging dually labeled transgenic *Hydra*

A major hurdle in recording *in toto* neural activity in a behaving *Hydra* is the faithful tracking of individual neurons over time as the animal moves. The initial transgenic *Hydra* created in our lab expressed GCaMP6s in all neurons, which allowed wide-field calcium imaging of whole animal neural activity while behaving^16^. However, each neuron had to be painstakingly tracked and analyzed manually. We then developed an automatic tracking algorithm (EMC^2^) that iteratively estimated the local deformation of the animal using spatial key points, and approximated the position of neurons when they were not firing and did not emit fluorescent signals^25^. This algorithm enabled us to track neurons with sufficient accuracy (>80%) over relatively long time-lapse sequences (∼1,000 frames at 10Hz). However, as tracking errors accumulate over time (when a neuron’s detection is associated to a wrong track, then all future detections of this neuron will also be erroneously associated), EMC^2^ was not sufficiently robust for long time-lapse sequences. To overcome this bottleneck, we developed a novel transgenic *Hydra* line that expresses two fluorescent indicators in its neurons: cytoplasmic GCaMP7s and nuclear tdTomato (Figure 2, see Methods section for more details). GCaMP7s is a genetically encoded calcium indicator (GECI) that allows visualization of the calcium dynamics of each neuron^31^ and tdTomato is a calcium insensitive molecule^32^ used for single cell labelling. tdTomato is a high brightness indicator with emission in the red end of the spectrum (581 nm) ideal for use with the green emission spectrum of GCaMP7s (511 nm) with minimal cross talk between the fluorophores. Our transgenic strain allows for the spatial tracking of the continuous signal from tdTomato, facilitating the tracking of individual neurons, and the extraction of GCaMP7s signal whenever neurons fire. The dually labeled transgenic line used in this protocol expresses these two fluorescent indicators in all cells derived from an initial interstitial cell. The interstitial cell lineage also produces germ cells, gland cells, and nematocytes^33–36^. The signal collected from these non-neuronal cells is later removed computationally (see section “Measuring the activity of individual neurons”).

**Figure 2.**
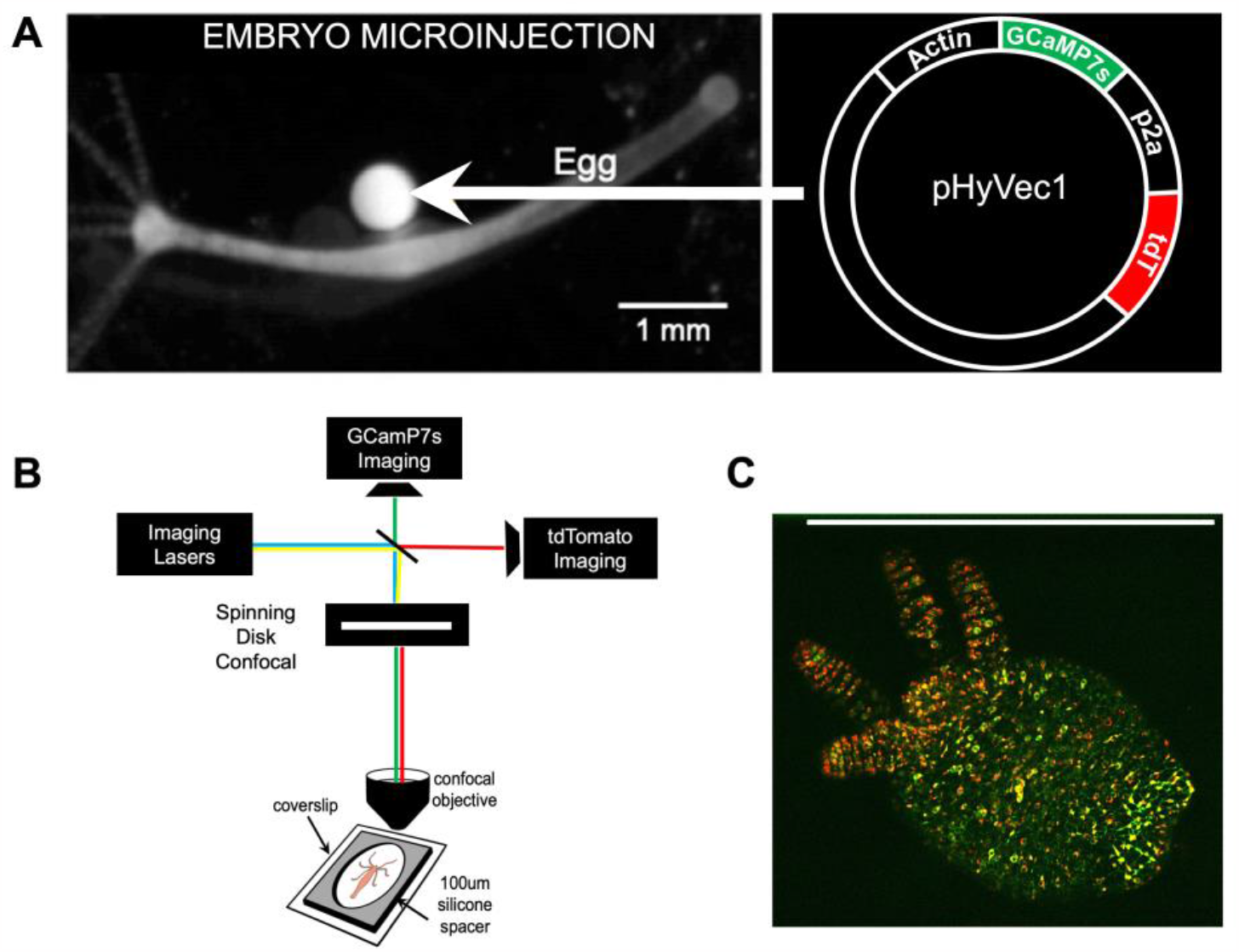
Generation of transgenic animals and imaging setup. (A) Transgenic animals were created by injecting the plasmid (right) into a fertilized *Hydra vulgaris* egg (left). Image on left modified with permission from^16^. (B) Dually labeled transgenic animals were placed in *Hydra* media in the center of a 100 *μm* silicon spacer sandwiched between two coverslips and imaged with a custom-built simultaneous two-color (488nm and 571nm) spinning disc confocal microscope. (C) Representative image of a dually labeled (nuclear tdTomato and cytoplasmic GCaMP7s) *Hydra* imaged with the simultaneous two-color spinning disc confocal microscope setup shown in (B). Scale bar is 1 mm.

The novel dually labeled transgenic *Hydra* was prepared for imaging as previously described^16^. In this mounted preparation, *Hydra* is placed in media between two coverslips separated by a 100 um spacer, creating an almost single-plane imaging object whilst allowing the animal to exhibit a variety of behaviors^37^. The animal is then imaged on a custom-built spinning disc confocal microscope that allows simultaneous two-color imaging of both GCaMP7s and tdTomato activity (see Methods). Confocal microscopy was adopted due to its superior performance over widefield, light sheet, and two-photon microscopy approaches in terms of its ability to perform simultaneous two-color recordings with sufficient spatiotemporal resolution and optimal signal-to-noise ratio. The fastest behaviors in *Hydra* occur at 5 Hz speed. Thus, to minimize motion blur in a rapidly contracting *Hydra*, images were acquired at 10 Hz. Recorded images were processed and adjusted for brightness and contrast using ImageJ analysis software.

### II-III. Segmenting and tracking individual neurons

Despite the calcium insensitiveness of nuclear tdTomato, the motion and deformation of the animal, the high density of neurons in some parts of the animal (peduncle and hypostome typically), and the potential occlusion or disappearance of neurons from the microscope focal plane due to axial motion of tissues, called for the development of a state-of-the-art pipeline for segmenting and tracking neurons’ nuclei.

With the exception of *C. elegans*, where neurons’ positions along time can be mapped and compared to an initial known atlas of positions after having corrected the deformation of the worm^38^, monitoring the calcium activity of individual neurons in other model animals requires the development of standard single-particle-tracking (SPT) that consists of two distinct steps: first, the fluorescent spots of individual particles (neurons) are automatically segmented, and the positions of detected neurons are extracted in each time frame of the movie. The second step of SPT algorithms is the robust linking of extracted positions into coherent tracks representing the trajectories of individual particles.

The first module of the TraSE-IN platform consists of a tracking pipeline, that we call ByoTrack (https://github.com/raphaelreme/byotrack) (Figure 3), which is a series of functions enabling the automatic detection of neurons’ nuclei and their tracking along the time-lapse sequence.

**Figure 3.**
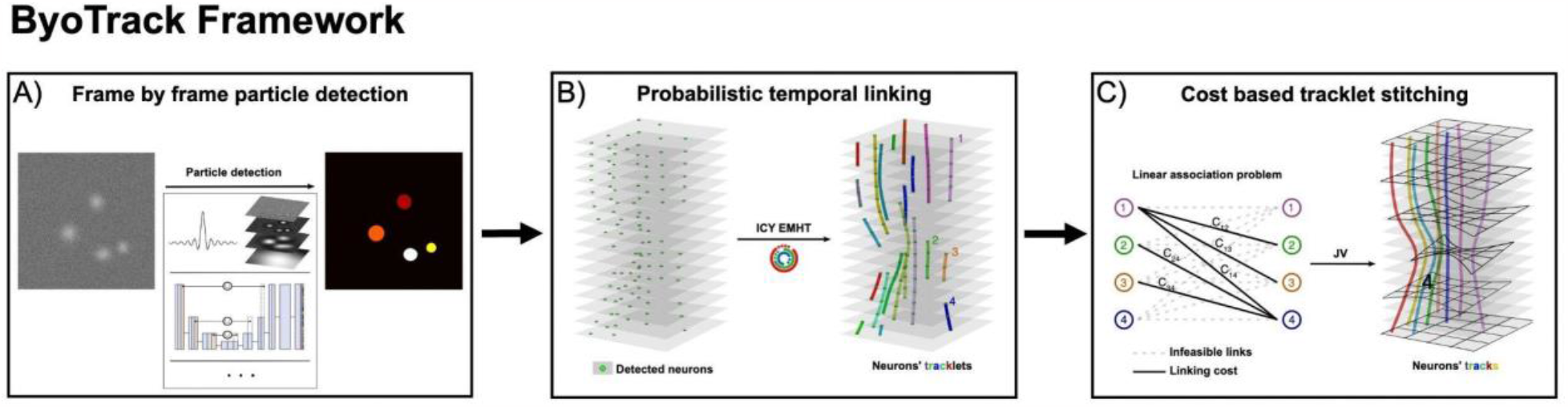
Tracking pipeline. The implemented tracking pipeline can be divided into three main steps: (A) the automatic detection of nuclei spots in tdTomato sequences using analytical methods such as wavelet transform and thresholding or deep learning methods such as StarDist detection, followed by (B) the robust linking of segmented nuclei in coherent tracklets with a state-of-the-art SPT method. Here, the probabilistic method enhanced Multiple-Hypothesis-Tracking (eMHT) method is called from the Icy software. (C) Finally, tracklets corresponding to the same neuron are stitched using a cost-based method. The cost here corresponds to the minimal distance between the forward- and backward-propagated positions of undetected nuclei. The optimal association between tracklets that minimizes the global cost of association is a linear association problem that we solved with the Jonker-Volgenant (JV) algorithm.

#### II. Segmentation of nuclei

In fluorescence microscopy, the objects of interest (neurons here) appear as spots brighter than the noisy background. When the signal-to-noise ratio (SNR) is low, objects’ spots can be hardly distinguishable. Therefore, elaborated detection algorithms have been developed over the years^39^. Most of the analytical detection methods consist of image denoising (e.g., Gaussian smoothing, wavelet denoising), signal enhancement (e.g., background subtraction with top-hats or h-dome), and signal thresholding. Analytical methods have proven to be robust and fast for moderate to high signal-to-noise ratios. However, they usually require tweaking a few user-defined parameters, and, more importantly, they usually fail at separating closely apposed spots.

Since their introduction, deep-learning (DL) approaches have demonstrated a much-increased accuracy and robustness for detecting cells in noisy and cluttered images. Among DL approaches, StarDist is particularly well suited for segmenting and separating roundish objects such as cell nuclei as it uses a star-convex representation of objects’ contours and a non-maximum-suppression (NMS) post-processing for separating touching cells^40^. However, DL approaches usually require manual annotation of data for model training.

In the first part of ByoTrack, we implemented different algorithms for neurons’ segmentation and detection (Figure 3A). In particular, we used an algorithm that executes the DL method StarDist, the training of the model being performed using the open-source Python code of StarDist^40^ . We also implemented an analytical method based on undecimated wavelet transform and statistical thresholding of the image ^41^. The wavelet thresholding approach has demonstrated a good performance on images with medium to low SNR^42^.

Using tdTomato images extracted from two-color time-lapse sequences we trained a StarDist model and compared its performance with the analytical wavelet method (Figure 4A). As manual annotation of cell nuclei can be tedious and time-consuming, we trained a StarDist model for an increasing number of tdTomato images (from 2 to 15, corresponding to a maximum of 9422 neurons’ nuclei, see Methods) and measured how model accuracy increased with the size of the training dataset (Figure 4B). In this manner, we could determine how to optimally balance model accuracy and training.

**Figure 4.**
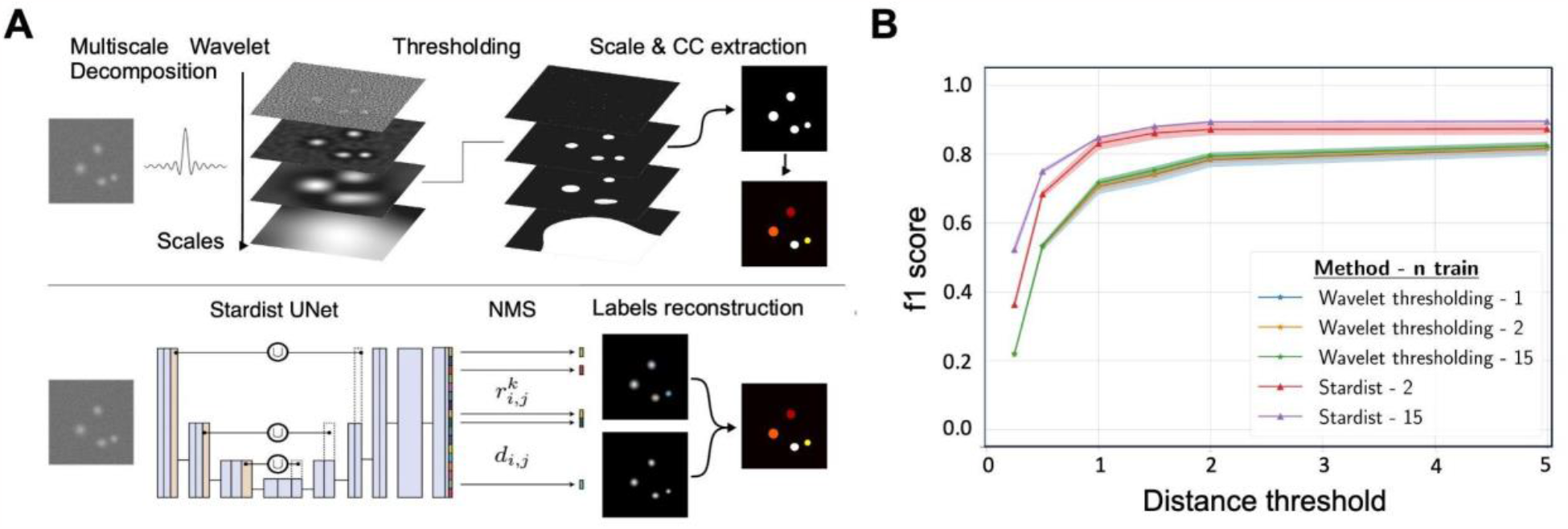
Deep-learning approach (StarDist) versus an analytical approach (wavelet transform) for segmenting nuclei in tdTomato images. (A) The two methods compared for detection. On top, the analytical wavelet transform approach consists of decomposing the image with multiscale *a-trou* wavelets, before applying a statistical threshold on wavelet coefficients to denoise the image and extract the significant fluorescent signals (connected components CC). Wavelet decomposition favors the extraction of spotty signals, which makes it well-suited for the segmentation of fluorescent roundish objects such as cell nuclei. On the bottom, StarDist deep learning (DL) segmentation consists of applying a trained U-net convolutional network, followed by a non-maximum suppression (NMS) to segment individual nuclei in the image. (B) We compared the accuracy (f1-score, mean ± standard error) of wavelet and DL segmentation algorithms applied to test images (*n* = 5 images) for different distance thresholds and number of images used for StarDist training (2 (red) or 15 (purple)), or for the calibration wavelet method’s parameters (scale and threshold: 1 (blue), 2 (orange) or 15 (green)).

Wavelet thresholding is specifically designed for the detection and localization of fluorescent spots, rather than for their accurate segmentation. Thus, the comparison metrics (f1-score) between detected spots was computed using a distance threshold *d* between the center-of-mass of detected objects, rather than the commonly used intersection-over-union (IOU) between segmented spots. In other words, we considered that a predicted spot’s detection was valid if its center-of-mass was at a distance less than *d* from the center-of-mass of a ground truth detection. Moreover, using the position of detected nuclei rather than their shape to assess the accuracy of the detection method is more meaningful for tracking, as we only use the position of each detected instance.

Using a distance threshold *d* = 1 pixel we obtained, for trained StarDist, a f1-score of 84.9% (number of predicted spots *n*_*pred*_ = 2946 with *t*_*p*_ = 2656 true positives and *f*_*p*_ = 290 false positives). The number of true instances (cell nuclei) was *n*_*true*_ = 3283 over 5 testing images, the number of false negatives being equal to *f*_*n*_ = 627 (Figure 4B). The hyper parameters of the analytical wavelet thresholding method (scale and threshold) were tuned (grid search) by maximizing the method accuracy over 1 to 15 annotated images. With the same distance threshold *d* = 1 pixel, we obtained a decreased f1-score of 72.0% (*n*_*pred*_ = 2747 *with t*_*p*_ = 2171 *and f*_*p*_ = 576). Number of false negatives was equal to *f*_*n*_ = 1112. Therefore, the false negatives and false positives were almost two-fold higher using an analytical wavelet-thresholding method compared to the trained DL method.

We highlight that StarDist performances were almost as good with only 2 annotated images (one image used for training, and one used for validation) compared to 15 annotated images for wavelet training. Therefore, we conclude that StarDist can be easily trained for other imaging set-ups or conditions and showed a much-increased performance compared to more standard, analytical detection algorithms.

#### III. Probabilistic tracking and tracklet stitching

To monitor the calcium activity of individual neurons, it is then necessary to link detected nuclei in the different time frames into coherent tracks. However, tracking is not trivial because of the errors made during the detection of particles (neurons) in each frame, the potential occlusion or undetectability of particles over several frames, and the difficult disentanglement of individual trajectories when particles are densely packed. Many elaborated algorithms have been developed in the last two decades^43^. These algorithms are either based on the global distance minimization (GDM) between the set of particles’ positions in successive time frames or use a probabilistic framework to compute the most likely trajectories linking the different positions given a model for particles’ motion. While requiring a higher computational load, the latter probabilistic framework has demonstrated an increased accuracy and robustness than GDM methods, especially in cluttered environments^44^. The second step of our tracking pipeline consists of linking neurons’ detections with a state-of-the-art probabilistic algorithm (eMHT, Figure 3B). This tracking algorithm is implemented in Icy, an open-source image analysis platform (https://icy.bioimageanalysis.org/)^45^. ByoTrack directly executes this step in Icy and gets the resulting tracks for post-processing and signal analysis.

Due to particles’ occlusion, moving out of focus, or intermittent detectability, standard tracking methods, such as eMHT, tend to produce multiple small tracks (“tracklets”) for the same particle, hindering the long-term identification of particles and, in the case of neurons, the monitoring of individual spiking activities. Indeed, when a particle is not detected over a few frames (more than ∼5 typically), SPT algorithms prematurely end tracks, and start a new track when the particle is re-detected.

To stitch the tracklets that represent the same neuron, we implemented the cost-based algorithm (EMC^2^)^25^ that infers the positions of undetected neurons from neighboring tracklets (Figure 3C). Briefly, when a neuron remains undetected over several frames, the forward position after the premature termination of the neuron’s tracklet is estimated using a thin-plate-spline interpolation between neighboring tracklets. On the other hand, when a new track begins, the backward position of the tracked neuron is also estimated. Then, to stitch tracklets and obtain entire neurons’ tracks, a cost matrix is computed between all the tracklets using the minimal distances between the forward and backward - estimated positions, and the corresponding linear association problem is solved with the Jonker-Volgenant algorithm^46^.

To measure the accuracy of our tracking pipeline on an experimental dataset, we tracked neurons’ nuclei using ByoTrack in *N* = 3 different animals (Figure 5). The rhythmic longitudinal contractions of *Hydra* can drastically hinder the robust tracking of individual neurons. Indeed, the speed of contraction motion can surpass the acquisition rate of the movie (10 Hz typically), blurring the images and making the detection and tracking of nuclei spots difficult. Thus, for each animal, we determined two time-windows of *T* = 1,000 frames where the animal contracts or not. For each of these six extracted movies (2 movies per animal, movies will be uploaded to Dryad), two independent operators chose randomly *n* = 15 individual neurons throughout the *Hydra* body (5 neurons in the animal’s peduncle (foot), central body, and hypostome (head)) and measured, for each neuron, the percentage of frames where it is correctly tracked. Obtained tracking accuracies are reported in Figure 5 (mean ± standard error), *n* = 30 neurons per localization (foot, body, and head) and animal state (contracting or not). We observe a very good accuracy (>90%) in all parts of the animal, except for the head of contracting animals where the accuracy drops to 70%. This local drop is due to the fast tentacle retraction and motion of the head of the animal during contraction cycles.

**Figure 5.**
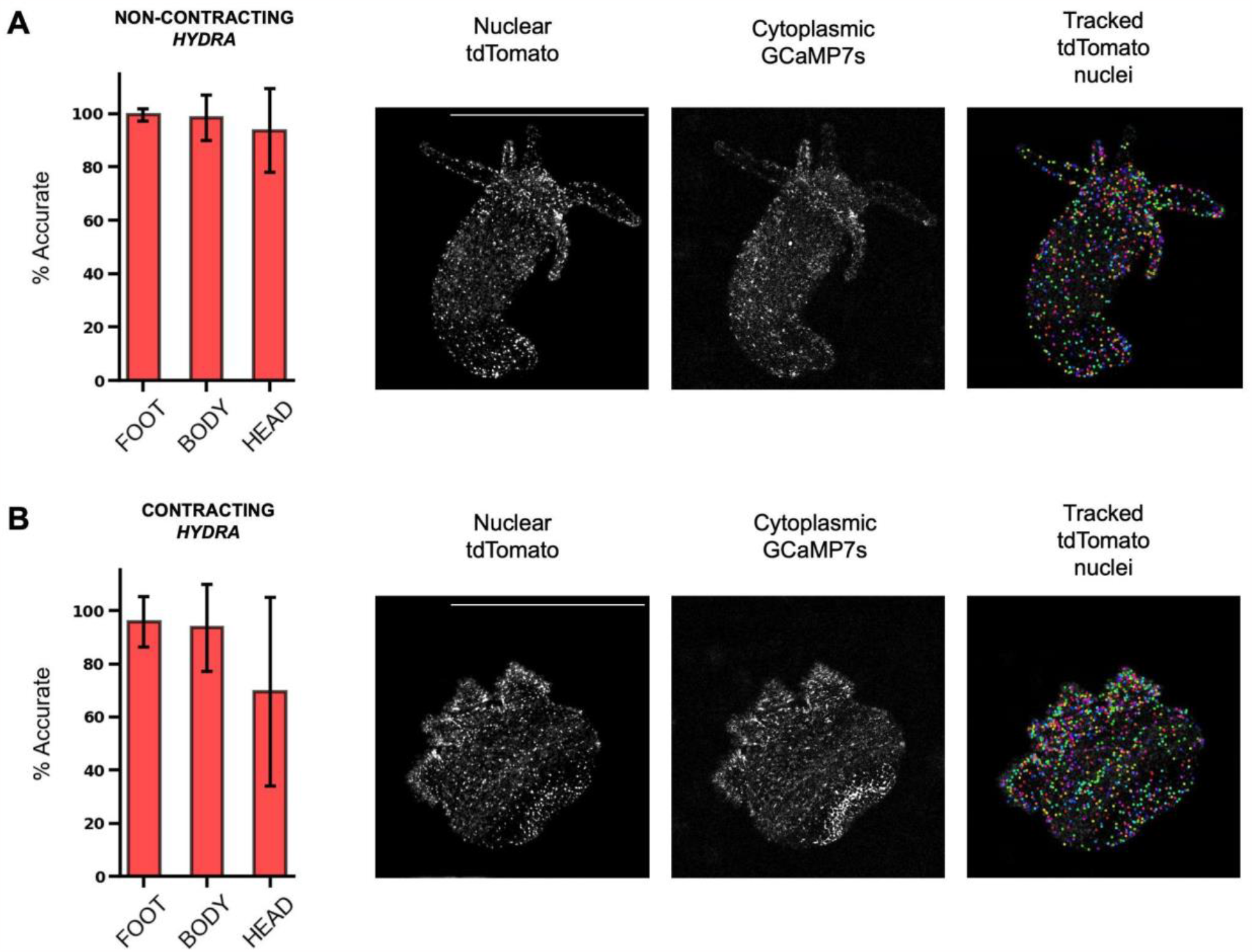
Accuracy of nuclei tracking in non-contracting and contracting *Hydra*. (A) Accuracy of tdTomato nuclei tracking in different parts of a non-contracting *Hydra* (left), with representative images of nuclear tdTomato, cytoplasmic GCaMP7s, and tracked tdTomato nuclei (right). (B) Accuracy of tdTomato nuclei tracking in different parts of a contracting *Hydra* (left), with representative images of nuclear tdTomato, cytoplasmic GCaMP7s, and tracked tdTomato nuclei (right) (mean ± std, *n* = 30 tracks). Scale bars are 1 mm.

Altogether, these results indicate that the ByoTrack pipeline is sufficiently versatile and robust to track individual neurons in highly deforming *Hydra*. In particular, the use of a DL segmentation algorithm should be preferred to more standard approaches because it does not require extensive training, while showing much improved accuracy. Except for the head of the animal that moves faster than the microscope’s acquisition rate during contraction phases, the combination of a state-of-the-art probabilistic tracking algorithm (eMHT) and cost-based tracklet stitching applied to DL-segmented nuclei, showed very good performance, paving the way to the monitoring of the calcium activity of individual neurons in behaving animals.

### IV-V. Measuring activity of individual neurons

#### IV. Linking calcium activity to tracked nuclei

The robust, long-term tracking of each neuron is a necessary prerequisite for the monitoring of calcium activity, but additional signal processing is required for the robust extraction of neurons’ spikes. First, even with perfect tracking of nuclei in the red channel, extracting the GCaMP7s signal is not as simple as mapping these tracks onto the video from the green channel. Indeed, optical and biological limitations cause misalignment between the two channels. Minor differences and imperfections in the optical path along with differing camera orientations cause each wavelength of light to be projected differently onto their respective cameras. Additionally, the indicators themselves are located in different parts of the cell, with tdTomato expressed in the cell’s nucleus and GCaMP7s being expressed in the cell’s cytoplasm. While some of this misalignment can be corrected with image processing software, the non-uniformity across the field of view is more complicated to correct. Moreover, multiple neurons often occur in close proximity, meaning the simple solution of taking the average fluorescence over a large region of interest (ROI) encompassing the nucleus will often include multiple neurons (Figure 6A). This can create confounding results where the signals from two neurons are superimposed (Figure 6B).

**Figure 6.**
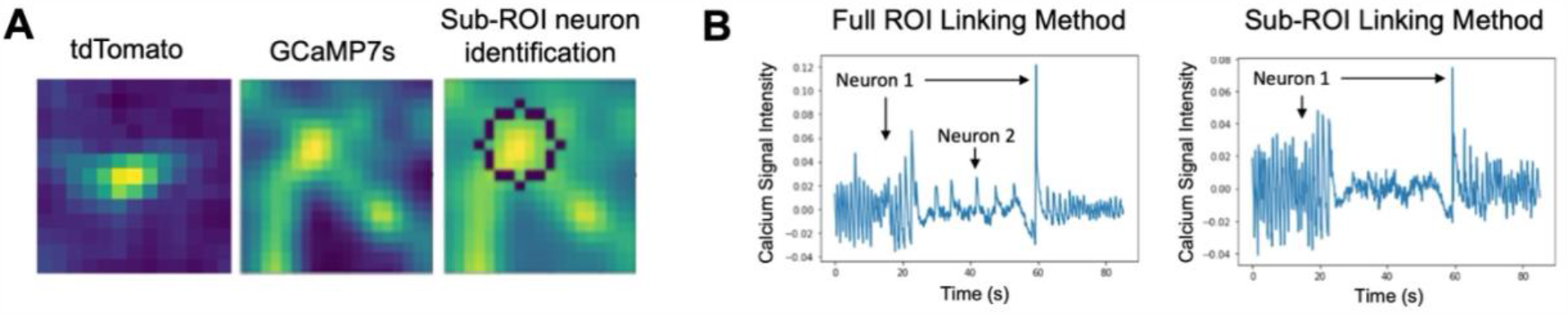
Linking tracked tdTomato nuclei to GCaMP7s neuronal cell bodies. (A) A small region of interest (ROI) is drawn around the center position of the tracked tdTomato nucleus (left). The brightest pixel nearest the center in the corresponding GCaMP7s image (middle) is used to detect the most likely neuron in the full ROI, around which a sub-ROI is drawn (right) and GCaMP7s signal extracted (see text for further details). (B) Example GCaMP7s activity when using the full ROI versus the sub-ROI linking method. In the full ROI method, the signal from two neurons in the same ROI is erroneously reported, whereas in the sub-ROI method only the signal from one neuron is extracted.

To handle this misalignment and extract robust single-neuron fluorescence information, a custom sub-ROI tracking method was developed (Figure 6) to identify and track the green-channel fluorescence in the area surrounding the red-tracked point. This algorithm starts from the pixel locations of tracked nuclei and extracts a small ROI around these points in the green channel. The algorithm leverages the fact that cells, even when not fluorescing, are brighter than the background. Within the small ROI, the total brightness, proximity to the center, and proximity to the previously tracked neuron are weighted and summed to give a prediction of the location of the neuron in the green channel (see Methods). This algorithm, implemented as a module on the TraSE-IN platform, provides reliable single neuron fluorescence data that can be further processed for spike extraction.

A second major hurdle to monitoring neuronal activity is the extraction of spikes from calcium fluorescence. Indeed, the fluorescent calcium traces are usually corrupted by background signal and noise, and the dynamics of conformational change of calcium indicators is much slower than actual voltage variations. For all these reasons, elaborate deconvolution methods have been developed over the years to robustly extract individual spikes ^26–28,47^. These algorithms show good performance on synthetic and experimental datasets but require an input fluorescent signal with good signal-to-noise ratio and no, or moderate, fluctuations of fluorescence baseline. To robustly extract neurons’ spikes from fluorescence intensity with these algorithms, we have therefore developed a series of pre-processing steps described hereafter (Figure 7).

**Figure 7.**
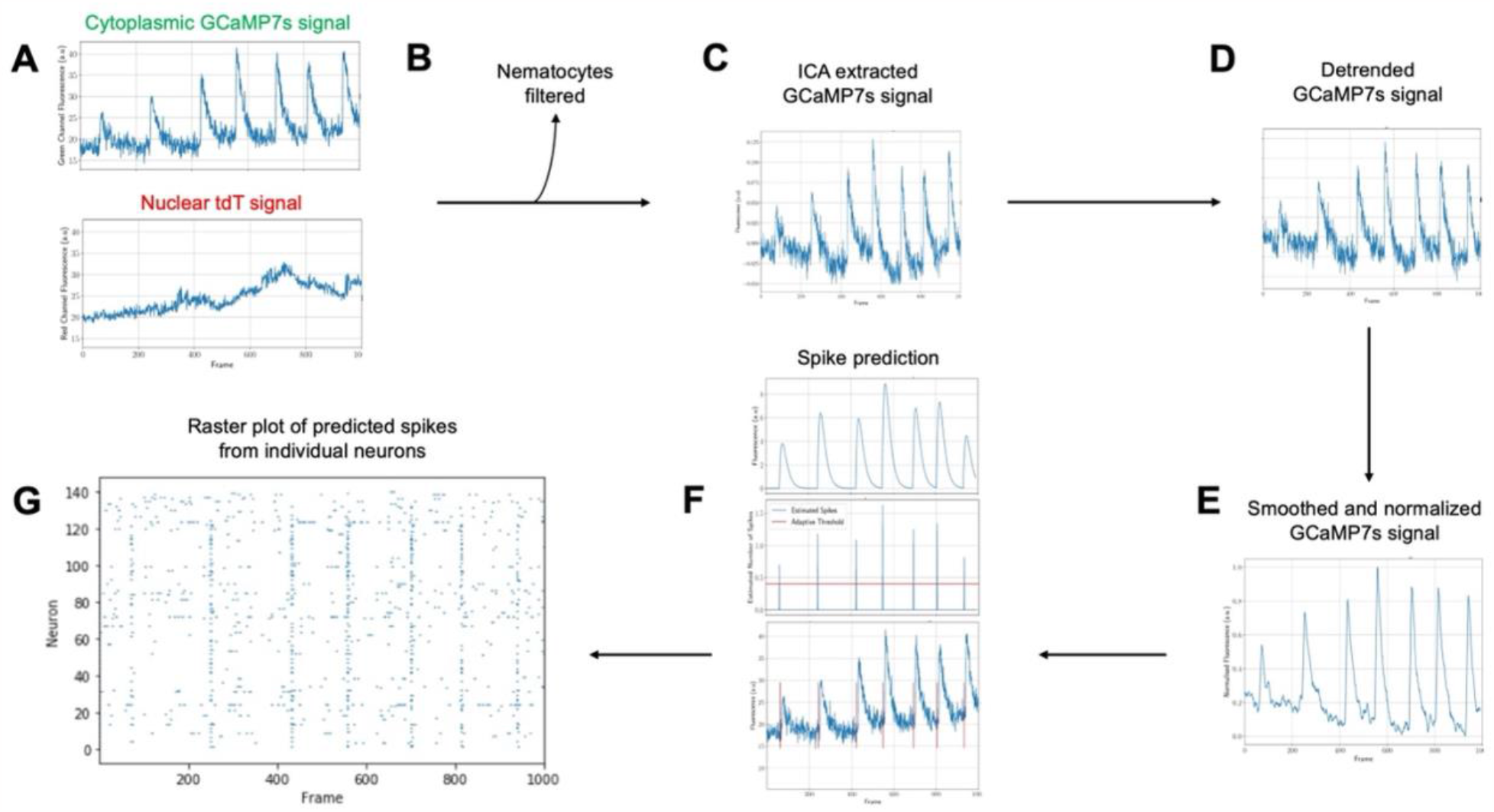
Schematic of signal processing and spike prediction steps. (A) Raw cytoplasmic GCaMP7s and nuclear tdTomato signals are extracted from individual neurons in each frame. (B) Nematocytes are filtered based on their correlation with Gaussian noise. (C) Independent component analysis (ICA) is used to extract the motion corrected GCaMP7s signal. (D) The motion corrected GCaMP7s signal is detrended with frequency filtering. (E) The detrended GCaMP7s signals from individual neurons are smoothed and (F) spikes are predicted using FOOPSI with a custom adaptive clustering and thresholding algorithm. (G) The final output is a raster plot of the predicted spikes from each individual neuron.

#### V. Signal processing and spike prediction

##### Removing Non-Neuronal Cells

Due to imperfect transgenic animals, where labeling occurs across all i-cell derived lines, non-neuronal cells express tdTomato and GCaMP7s. Most of these cells do not display calcium activity so are simple to remove or ignore. Nematocyles, the stinging cells unique to cnidarians, are an exception. These cells produce very bright calcium signals and will be incorrectly analyzed by spike extraction methods if not removed.

Filtering nematocytes can be achieved either via their morphology (they are larger than neuronal cells) or their signal. We chose to filter nematocytes based on their signal due to its comparative simplicity. As these cells do not spike like neurons, their calcium signals are almost entirely noise. Leveraging this fact allows filtering of nematocytes (Figure 7B) based on their correlation with Gaussian noise (see Methods)

##### Motion Artefact Removal

Motion artifacts occur when *Hydra* moves axially during imaging. Due to the narrow focal plane of confocal microscopy, and occlusion of light from outside this focal plane, any motion in Z will result in a significant change in intensity, and as *Hydra* is especially deformable, this type of motion occurs somewhat often in behaving animals. Additionally, due to the non-uniformity of *Hydra* motion, these artifacts do not apply to the entire frame at once, making some general motion correction algorithms useless.

Motion artifacts are notoriously challenging to correct algorithmically as they are highly sporadic, and the intensity and duration of their effect varies at random. However, we can leverage our simultaneous two-color imaging to provide some measure of correction. Since motion occurs in both the red and green channels identically, any large signal occurring in both channels is likely to be a motion artifact and we can use a variety of methods to remove it.

The standard method for removing motion artifacts in a two-color experiment is ratiometric correction. However, this method can lead to erroneous spikes being recorded: when both signals approach low values, the ratio between them can be very large even if the absolute difference is small. Therefore, ratiometrically corrected signals can show a large increase in intensity when both signals are actually decreasing in intensity.

Instead, we used independent component analysis (ICA) (Figure 7C) to separate the motion artifacts from changes in fluorescence due to calcium dynamics^48^. This method allows source separation by directly looking for common signals between the channels. ICA assumes that the only signals present in both the red and green channels are motion artifacts and calcium signals and seeks to find two component signals that can be recombined in differing proportions to reconstruct the original red and green channels. In practice, this method produces two signals: one mostly composed of motion signal, and the other composed of calcium signal. Noise is distributed equally between the outputs. As ICA is a stochastic algorithm, our implementation runs multiple times and selects the best resulting signals (see Methods). The calcium signal is identified from the outputs via its higher correlation with the original green channel signal. We found this method to produce significantly better results than ratiometric correction.

##### Detrending and Smoothing

Although ICA can reduce artifacts that occur in both channels identically, it cannot completely attenuate them, nor can it process artifacts that occur differently in each channel such as photobleaching, which occurs at different rates in tdTomato and GCaMP7s. Additionally, spike identification algorithms are fragile. Baseline drift or abundant noise can drastically hinder their performance and cause them to miss or to incorrectly predict spikes, making downstream analysis far less reliable.

To improve the performance of our spike extraction, two additional preprocessing steps were performed: detrending to remove baseline drift (Figure 7D) and smoothing (Figure 7E) to clean the signals by reducing noise. Detrending was applied via frequency filtering. We deleted from each signal the frequencies smaller than a threshold tuned with user feedback. Smoothing was applied using a rolling average filter and was again tuned with user feedback to allow for optimal reduction of noise without removal of signal (see Methods).

##### Calcium Signal and Spike-Time Extraction

After filtering and preprocessing the extracted fluorescence signals, we utilized the FOOPSI spike extraction method (Figure 7F) implemented in the CaImAn Python package^30^. This package is intended for use with mouse two photon calcium imaging data. However, since our data has low background noise and uses the same indicator used in mouse studies, we found this method was reliably able to extract spikes from our processed data with only minor tuning.

FOOPSI fits an autoregressive model to the provided signal and, using an underlying model of the expected calcium signal dynamics from GCaMP indicators, can predict the underlying fluorescence of GCaMP7s while ignoring the accompanying noise. In addition to extracting fluorescence, the FOOPSI algorithm provides spike times predicting the underlying electrical activity of the neuron from its calcium activity.

Spike time information from FOOPSI is provided as a probability of a spike having occurred at any given point in time. To extract robust spike times, a custom clustering and thresholding algorithm was developed (see Methods). The final output of the signal processing module is a raster plot (Figure 7G) with predicted spikes for all tracked individual neurons.

## Methods

### Generation of transgenic *Hydra*

Following the method described by Juliano et al.^49^, a transgenic *Hydra vulgaris* (strain AEP) line was established to express GCaMP7s in the cytosol and tdTomato in the nucleus of cells derived from the interstitial stem cell lineage. The plasmid (Addgene catalog no. 102558)^16^ was modified by replacing the actin promoter with the EF1-alpha promoter and inserting DsRed downstream of the GCaMP6s sequence. To ensure the nuclear localization of DsRed, a P2A self-cleaving peptide sequence containing a nuclear localizing signal (cccaagaagaagaggaaggtg) was inserted between GCaMP6s and DsRed. The nuclear localization of DsRed was confirmed by electroplating the plasmid into *Hydra*. To enhance fluorescence intensity, GCaMP6s was replaced with jGCaMP7s and DsRed was replaced with tdTomato. Finally, for microinjections, the EF1-alpha promoter was replaced with the Actin promoter. All fluorescent reporter gene sequences were codon optimized specifically for *Hydra*. Standard embryo microinjection of this plasmid was performed and transgenic hatchlings expressing cytoplasmic GCaMP7s and nuclear tdTomato in the interstitial cell lineage were isolated. Transgenic animals were bred until all neurons were expressing both cytoplasmic GCaMP7s and nuclear tdTomato. Transgenic *Hydra* were cultured using standard methods^50^ in *Hydra* medium (1mM calcium chloride dehydrate, 0.33mM magnesium sulfate anhydrous, 0.5mM sodium bicarbonate, 0.03mM potassium chloride) at 18°C on a 12 hr light/12 hr dark cycle. They were fed freshly hatched *Artemia* <u>nauplii</u> twice per week.

### Imaging

Dual-labeled transgenic *Hydra* were prepared for imaging as described^16^. Imaging was performed using a custom dual-channel spinning disc confocal microscope (Solamere Yokogawa CSU-X1) with a sCMOS camera for each channel (Teledyne-Photometrics Prime-BSI). Samples were simultaneously illuminated with both 488 nm and 561 nm lasers (Coherent OBIS). Emission light was split with a dichroic mirror sending green light to one camera and red light to the other. Cameras were aligned with pinholes and images were registered during processing. Images were captured with a frame rate of 10 frames per second using a 6X objective (Navitar HR Plan Apo 6X/0.3) and Micro-Manager software.

### Segmentation and tracking of nuclei (ByoTrack)

#### Segmentation of neuronal nuclei

To learn a segmentation model and validate our experiments, we sampled 20 tdTomato images of *Hydra* on various positions of the animal (e.g., contracted, elongated). We manually annotated these images using the ImageJ draw tool resulting in 12,705 segmented neurons (∼600/image).

To measure the segmentation performance, 5 random images were kept as a testing dataset. To account for performance variability, we repeated this process 5 times with different seeds, resulting in different training and testing images. We measured the performance with each seed and report the results as (mean ± std).

We used standard instance segmentation metrics (precision, recall, f1-score). But we did not base these metrics on IOU association, where a predicted instance is considered to be a true positive if it overlaps with a ground truth on a sufficiently large area. We rather based these metrics on distance association, where a predicted instance (spot) is considered valid if its mass-center is sufficiently close to the mass-center of a ground truth instance. This has two benefits: first, it is more meaningful for tracking as we only use the position of each detected instance. Second, it allows fair comparison with detection methods that do not try to produce a faithful segmentation or bounding box of the instances like the wavelet thresholding method.

The remaining 15 images were used to train and tune all the parameters of the segmentation model. We also measured the impact of the training data quantity by training the model with fewer images, from 1 to 15.

We compared two detection methods: an analytical method based on wavelet decomposition and thresholding, and a deep learning approach, StarDist.

#### Calibrating wavelet thresholding method

The method has only two hyperparameters: the scale of the spots and the noise threshold. Both can be easily tuned by hand on a single image. Nonetheless, to be fully autonomous, the system performs a grid search on the training images and chooses the best performing ones. No further training is required. Moreover, to reduce the number of false positives, the neurons with less than 5 pixels area were filtered out.

#### Training DL method (StarDist)

The deep neural network must be trained and the hyperparameters tuned. We therefore split the training images into training and validation as most standard deep learning approaches do. 20% of the training images were used as validation images. With less than 5 training images, one image was kept for validation. Therefore, the procedure that we show here requires at least 2 training images, one to train the weights of the network, the other to validate the hyper-parameters.

The official implementation of StarDist (https://github.com/stardist/stardist) was used to train, validate, and evaluate the performances with our dataset. For tracking, we provide a wrapper to perform the detection process using a trained StarDist model (https://partage.imt.fr/index.php/s/npwHJHZebxqGMPi).

#### Generating tracklets with probabilistic eMHT

The eMHT algorithm is implemented on the open-source imaging platform Icy (plugin Spot Tracking), which is coded in Java language. Instead of translating all the code into Python, we decided to directly call the Spot Tracking plugin in Icy in a headless mode from the TraSE-IN platform. To ensure compatibility with Icy, we implemented different functions for inputs (detections) and outputs (tracks) wrapping.

We believe that calling implemented tracking solutions in major imaging platforms such as Icy and ImageJ is an efficient and sustainable solution because the main imaging platforms are open-source, the tracking plugins (e.g., Spot tracking in Icy, Trackmate in ImageJ) are regularly updated by developers, wrapping input and output variables in TraSE-IN is much less tedious and prone to implementation errors than translating all the Java code of tracking methods in Python, and TraSE-IN will easily integrate new tracking solutions developed on other platforms, if needed.

#### Stitching tracklets

To stitch tracklets obtained with eMHT tracking, we implemented a cost-based algorithm. Briefly, a cost between all the tracklets is computed, and stitched tracklets correspond to the minimal global cost of all tracklets’ linking. Computing the minimum global cost corresponds to a linear assignment problem that we solved using the Jonker-Volgenant algorithm^46^. The cost between tracklets was computed following the EMC^2^ algorithm^25^. Input parameters of the stitching function (byotrack.implementation.refiner.stitching.emc2) are the smoothness *α* of the Thin Plate Spline interpolation and the non-linking cost *η* (paid for unlinked tracklets). In our tracking scenario (Figure 4C), we set *α* = 10 to provide enough regularization to be robust to eMHT linking errors and set non-linking cost *η* = 5 pixels.

### Linking calcium activity to tracked nuclei

To solve the misalignment between nuclei and calcium signal, we implemented a sub-ROI tracking method. For each nucleus, we extracted a 25×25 pixels ROI centered on each tracked nucleus. Then, to find neuronal candidates in the GCaMP7s channel, we iteratively computed *n* local maxima within each ROI (*n* = 5 typically) and identified the most probable calcium signal. For this, we first applied a gaussian smoothing (*σ* = 1 pixel) and first identified the global maximum in the ROI. We then deleted the pixels in the neighborhoods of this maximum and reiterated the process *n* times. To bias extracted maxima towards the nucleus position (and avoid selecting an outlier GcaMP7s signal) we weighted the ROI intensities with Gaussian prior centered on the nucleus position (*σ* = 5 pixels).

In the first frame, the selected position of the calcium spot is the global maximum of the ROI (after having applied a Gaussian prior to bias intensities towards the nucleus position). For the following frames, we selected the closest maximum to the previous calcium position and updated the calcium position with an exponential moving average to attenuate the potential consequence of a wrong association. Finally, we extracted the calcium intensity for each neuron as the mean intensity within the 5 pixels radius circle centered on the tracked calcium position.

### Signal processing

#### Removing Non-Neuronal Cells

We compared extracted calcium signals with a normal distribution using the scipy.stats.normaltest method. Any signal that can be rejected (different than normal) with p-value greater than a user-defined threshold was kept, and the others removed. To improve the filtering step, an interactive feedback step was implemented allowing users to correct the filtering or find the best threshold until most nematocytes are removed without erroneously removing any neurons.

#### Motion Artefact Removal

We used the tdTomato intensity (extracted as the mean intensity in a 5 pixels radius circle centered on the nucleus position in the red channel) as a control signal. ICA between the control signal and the raw calcium signal was used to remove motion artefacts^48^. ICA relies on scikit-learn FastICA method^51^ (we used a tolerance of 0.5). As the ICA algorithm is stochastic, we ran it several times and selected the best outcome as the one with the minimum correlation with the control signal.

#### Detrending & Smoothing

To remove the remaining baseline in the calcium signal, we applied frequency filtering using a 5^th^ order Butterworth filter and a critical period of 100 frames. In addition, we smoothed the signal using a rolling average of size 5. Both the average size and critical period can be adapted with user feedback.

#### Calcium Signal and Spike-Time Extraction

We used the constrained FOOPSI implementation in the CaImAn Python library (OASIS method of order 2)^30^. In *Hydra*, calcium spikes correspond to single action potentials^16^. Therefore, we added a clustering step to aggregate the action potentials (electrical spikes) predicted with the previous deconvolution FOOPSI algorithm (less than 5 frames away). The clustering consists of first temporal Gaussian blurring (*σ* = 5 frames) of the estimated electrical spikes, followed by local maxima extraction. Finally, we filtered out electrical spikes with low probability/intensity using a user-defined threshold to reduce the number of erroneous predictions. Since these computed probabilities/intensities vary wildly between different signals, using a fixed threshold would not work across different signals. To rectify this, an adaptive threshold defined as the maximum probability minus 2 standard deviations was used. This method provided robust results across all cells, allowing spike extraction to be applied simultaneously across an entire dataset with minimal user input.

## Discussion

Here, we present TraSE-IN, an open access end-to-end protocol for single-neuron resolution calcium imaging in entire behaving animals. To our knowledge, this pipeline is the first to allow automatic detection, tracking, signal extraction, and spike prediction from all neurons in a highly deformable organism such as *Hydra*.

Hardware improvements implemented in this pipeline include a novel dually labeled transgenic *Hydra* that is imaged with simultaneous two-color spinning disc confocal microscopy to allow high spatiotemporal resolution imaging of the entire animal in the field of view while behaving. The novel transgenic *Hydra* presented here is labeled with a calcium-insensitive fluorophore (tdTomato) in the nucleus and calcium-sensitive fluorophore (GCaMP7s) in the cytoplasm of each neuron. The presence of the calcium-insensitive fluorophore in the nucleus of each neuron allows visualization of each neuron even while it is inactive and significantly increases accuracy of tracking individual neurons over time. The high-resolution spinning disc confocal imaging also increases the accuracy of tracking individual neurons longitudinally because they are easier to detect in each frame due to the high spatial resolution, and easier to track frame-to-frame due to the high temporal resolution. Software improvements include more accurate segmentation of neurons in each frame using a machine learning algorithm (StarDist^40^) specifically trained to detect *Hydra* nuclei, an accurate single neuron tracking protocol using a state-of-the-art probabilistic algorithm (eMHT^44^) that is directly executed on the native open-source platform Icy^45^, tracklet stitching^25^, and, finally, a user-friendly Python pipeline for signal extraction, signal processing, and spike prediction for all neurons in an entire behaving animal.

While the development of this pipeline is a major step forward towards the goal of automatic detection, tracking, signal extraction, and spike prediction from single neurons in whole behaving animals over longer timescales, multiple areas for future improvements remain. First, although the novel dually labeled transgenic *Hydra* line allows visualization of all neurons even while inactive, the label is not specific to neurons. The label is expressed in the interstitial cell lineage of the animal, which is a stem cell lineage that makes four cell types: neurons, nematocytes, gland cells, and germ cells^33–36^. Thus, all these cell types express the label in the transgenic animals. In the current pipeline, the non-neural cell types are filtered out in the signal processing step of the pipeline. Instead of filtering out these different cell types during signal processing, it would be ideal to generate an animal with neuron-specific labeling using neuron-specific promoters^17^. In addition to more specific labeling of neurons in animals, it would be ideal to use different optical approaches that allow three-dimensional volumetric imaging of the entire animal while behaving. Here, *Hydra* is only imaged in one two-dimensional plane, which captures most of its neurons, but this method results in some neurons coming in and out of the imaging plane, so some neurons are lost over time. This is especially problematic for the tentacles, which move much more than the body column of the animal. At present, current volumetric approaches are too slow to allow high spatiotemporal resolution imaging of an entire rapidly moving organism^24^. However, as fluorescent labels and optics improve, shorter exposure times and higher frame rates may be possible to allow whole animal volumetric imaging while *Hydra* is behaving.

In addition to hardware improvements, multiple improvements can also be implemented on the software side. In terms of segmentation, even if Stardist provides accurate detection with relatively few manual annotations (∼ 2 image patches for a total of ∼ 1,000 segmented nuclei), transfer learning from pre-annotated public datasets would be a very valuable resource for the segmentation of cells in various imaging set-ups. In terms of tracking, we designed TraSE-IN so it does not code for a specific method, but rather executes state-of-the-art tracking algorithms directly from their native platform. In the present version, TraSE-IN is executing the eMHT probabilistic algorithm in Icy (https://icy.bioimageanalysis.org/), but future versions of the platform could implement other well-established tracking methods such as TrackMate in ImageJ^52^. For the tracklet stitching, the cost between tracklets corresponds to the minimal distance between the predicted forward- and backward-propagated positions of tracklets’ ending- and starting-points, respectively^25^. This cost could also include visual features of tracklets, which should further improve the accuracy and robustness of tracklets’ stitching, as recently shown^53^. In terms of signal extraction, many putative improvements reside in spike extraction. Indeed, the robust estimation of spiking times from noisy fluorescence traces is an active field of research, and recent developments in Bayesian analysis^26,47^ might improve the accuracy of spike estimates compared with previous constrained deconvolution methods^27,29,54^.

In sum, we have created a robust, adaptative end-to-end pipeline that allows single-neuron resolution imaging of an entire behaving *Hydra* followed by automatic detection, tracking, signal extraction, signal processing, and spike prediction of all individual neurons. While room for future improvement remains, this new tool takes us one step closer towards the goal of automatically extracting every spike from every neuron in a behaving animal over longer timescales. Such data will allow visualization and analysis of the emergent properties of an entire nervous system under different conditions, making it possible to eventually crack the neural code of a whole animal.

## Authorship Contributions

A.H. injected and screened the transgenic *Hydra* line, developed the custom imaging method and acquired all images, trained the StarDist model, and guided development of the platform. R.R. and N.T. conceived and implemented the TraSE-IN platform. W.Y. initiated a dual-color tracking project by constructing and validating the reporter plasmid using electroporation and injections. A.H, T.L, R.R., and N.T. analyzed data. All authors contributed to project conception, design, and manuscript writing. J-C.O-M, T.L. and R.Y. assembled the team, directed the project, and secured equipment, resources and funding.

## Data Availability Statement

Source code and documentation for the plugin are available at https://github.com/raphaelreme/trase-in. Movies will be available on the Dryad platform.

